# Complete genome sequence and annotation of the laboratory reference strain *Shigella flexneri* serotype 5a M90T and genome-wide transcriptional start site determination

**DOI:** 10.1101/595066

**Authors:** Ramón Cervantes-Rivera, Sophie Tronnet, Andrea Puhar

## Abstract

**Background:** *Shigella* is a Gram-negative facultative intracellular bacterium that causes bacillary dysentery in humans. *Shigella* invades cells of the colonic mucosa owing to its virulence plasmid-encoded Type 3 Secretion System (T3SS), and multiplies in the target cell cytosol. Although the laboratory reference strain *S. flexneri* serotype 5a M90T has been extensively used to understand the molecular mechanisms of pathogenesis, its complete genome sequence is not available, thereby greatly limiting studies employing high-throughput sequencing and systems biology approaches.

**Results:** We have sequenced, assembled, annotated and manually curated the full genome of *S. flexneri* 5a M90T. This yielded two complete circular contigs, the chromosome and the virulence plasmid (pWR100). To obtain the genome sequence, we have employed long-read PacBio DNA sequencing followed by polishing with Illumina RNA-seq data. This provides a new pipeline to prepare gapless, highly accurate genome sequences. Furthermore, we have performed genome-wide analysis of transcriptional start sites and determined the length of 5’ untranslated regions (5’-UTRs) at typical culture conditions for the inoculum of *in vitro* infection experiments. We identified 6,723 primary TSS (pTSS) and 7,328 secondary TSS (sTSS). The *S. flexneri* 5a M90T annotated genome sequence and the transcriptional start sites are integrated into RegulonDB (http://regulondb.ccg.unam.mx) and RSAT (http://embnet.ccg.unam.mx/rsat/) to use its analysis tools in *S. flexneri* 5a M90T genome.

**Conclusions:** We provide the first complete genome for *S. flexneri* serotype 5a, specifically the laboratory reference strain M90T. Our work opens the possibility of employing *S. flexneri* M90T in high-quality systems biology studies such as transcriptomic and differential expression analyses or in genome evolution studies. Moreover, the catalogue of TSS that we report here can be used in molecular pathogenesis studies as a resource to know which genes are transcribed before infection of host cells. The genome sequence, together with the analysis of transcriptional start sites, is also a valuable tool for precise genetic manipulation of *S. flexneri* 5a M90T. The hybrid pipeline that we report here combining genome sequencing with long-reads technology and polishing with RNAseq data defines a powerful strategy for genome assembly, polishing and annotation in any type of organism.

## Background

*Shigella* is an enteroinvasive Gram-negative bacterium that causes shigellosis or bacillary dysentery in humans. *Shigella* is responsible for significant morbidity and mortality, particularly in young children and immunocompromised adults[1, 2]. In 2010, around 188 million cases of shigellosis occurred globally, including 62.3 million cases in children younger than 5 years[3–5]. A vast majority of the disease burden due to *Shigella* spp. can be attributed to *S. flexneri* in the developing world and to *S. sonnei* in more industrialized regions[1].

*S. flexneri* has a low infection dose of only 10 to 100 bacteria[6]. *Shigella* causes disease by invading the colonic mucosa, resulting in an intense acute inflammatory response. The bacterium spreads via the fecal-oral route upon ingestion of contaminated food or water and also via person-to-person contact[7].

*S. flexneri* 5a M90T, along with *S. flexneri 2a*, is the most commonly employed laboratory reference strain for *s. flexneri*[8]. Indeed, the majority of our knowledge of the molecular mechanisms of *Shigella* pathogenesis has been obtained using *S. flexneri* M90T as a model. The genome of this strain is composed of a circular chromosome and a megaplasmid (virulence plasmid), called pWR100[9]. The pathogenesis of *Shigella* spp. strictly depends on the virulence plasmid, which encodes several factors that are essential for invasion of host defenses[10].

Due to its prime importance, the virulence plasmid was one of the first genomic elements to be sequenced, at least partially, in *S. flexneri* 5a M90T; a major breakthrough at the time[10, 11]. The virulence plasmid, later renamed pWR501[10, 11], was sequenced using Applied Biosystems ABI377[11]. This now-obsolete technology required nebulization and subsequent size fractionation (in the range of 0.7 to 2.0 kb) of DNA by agarose gel electrophoresis and cloning into cosmids for sequencing[11]. This protocol increased the risk of introducing mutations or losing fragments, which can now be ameliorated by the use of NGS technology such as PacBio/SMRT long-read sequencing[12], which is cloning- and PCR-free.

So far, chromosomally encoded genes have received little attention, as most *Shigella* research has focused on the plasmid-encoded virulence genes. However, some of the genes encoded on the chromosome may play an important role in *Shigella* pathogenesis[13]. The *S. flexneri* 5a M90T chromosome has been sequenced and assembled earlier[14], but this sequence is not complete, it is only reported as a genome scaffold with many gaps. Moreover, the sequence assembly and annotation was based on another *S. flexneri* strain, *S. flexneri* serotype 5b 8401[15]. To better understand the pathogenic mechanisms and to identify the genetic elements that are involved in pathogenicity and its regulation, it is essential to have a fully sequenced and annotated genome.

Transcriptomic analysis has been increasingly employed to dissect the molecular mechanisms of host-pathogen interactions for a wide range of bacteria[16–19]. However, only few studies employing RNA-seq have been carried out in *Shigella*[13, 20, 21]. The lack of a *S. flexneri* 5a M90T high-quality genome for transcriptome data analysis is a hinderance, leading to poor reads alignment in our experience. Thus, the availability of the annotated full genome of *S. flexneri* serotype 5a strain M90T paves the way to use this model organism for molecular pathogenesis studies by transcriptome analysis. Taken together, in spite of the wealth of molecular pathogenesis data obtained with *S. flexneri* 5a M90T, we are still in need of a complete and high-quality genome sequence for this strain[21].

Genes in prokaryotic cells can have more than one transcriptional start site (TSS). Typically, transcription starts at position 20/-40 from the first translatable codon[22]. However, it is already known that in many bacteria the transcriptional start site is variable, depending on the environment. Further, it is also known that TSSs vary depending on how bacteria respond to a specific stimulus[23]. Knowing the operon and gene structure is essential to understand gene expression and regulation. Hence, the determination of the transcriptional start site is one of the first steps in understanding the molecular mechanisms that are implicated in the regulation of a gene.

Primary transcripts of prokaryotes carry a triphosphate at their 5’-ends. In contrast, processed or degraded RNAs only carry a monophosphate at their 5’-ends[24]. The differential RNA-seq (dRNA-seq) approach used here exploits the properties of a 5’-monophosphate-dependent exonuclease (TEX) to selectively degrade processed transcripts, thereby enriching for unprocessed RNA species carrying a native 5’-triphosphate[24]. TSSs can then be identified by comparing TEX-treated and untreated RNA-seq libraries, where TSSs appear as localized maxima in coverage enriched upon TEX-treatment[16].

Here we present the full, high-quality, and annotated genome of *S. flexneri* 5a M90T. Furthermore, we identified the genes that are expressed during mid-exponential growth in TSB, the typical condition used for *in vitro* infections with *Shigella*. In addition, we determined the active transcriptional start sites during mid-exponential growth in TSB and the length of 5’-UTR regions.

## Results

### Complete and gapless genome assembly of *S. flexneri* 5a M90T

To determine the genome sequence of *S. flexneri* serotype 5a strain M90T whole-genome sequencing was conducted with 3-cell sequencing in a PacBio single-molecule real-time (SMRT) sequencing system[12]. This generated a raw output of 93,316 subreads with mean length of 8,387 bp and the longest read of 12,275 bp. The sequences totaled 782,710,041 bp, which corresponds to ∼157-fold genome coverage. This coverage is high enough to avoid any possible sequencing error.

Genome assembly was carried out with Canu/1.7[25], feeding PacBio raw data. This assembly generated two contigs without any gap and suggested circular replicons. For the larger contig, the output from Canu retained 14,193 reads of 5,938 bp average read length, with a total contig length of 4,596,714 bp (Fig. 1a), indicating that this contig corresponds to the chromosomal replicon. For the smaller contig, Canu retained 1491 reads of 5,938 bp average read length, with a total length of 232,195 bp (Fig. 1b). The small size of this replicon suggested that it corresponds to the virulence plasmid. These two replicons roughly correspond to the expected size for the chromosome and virulence plasmid of *S. flexneri* 5a M90T, in accordance with previous reports[10, 11, 14].

**Figure 1.**
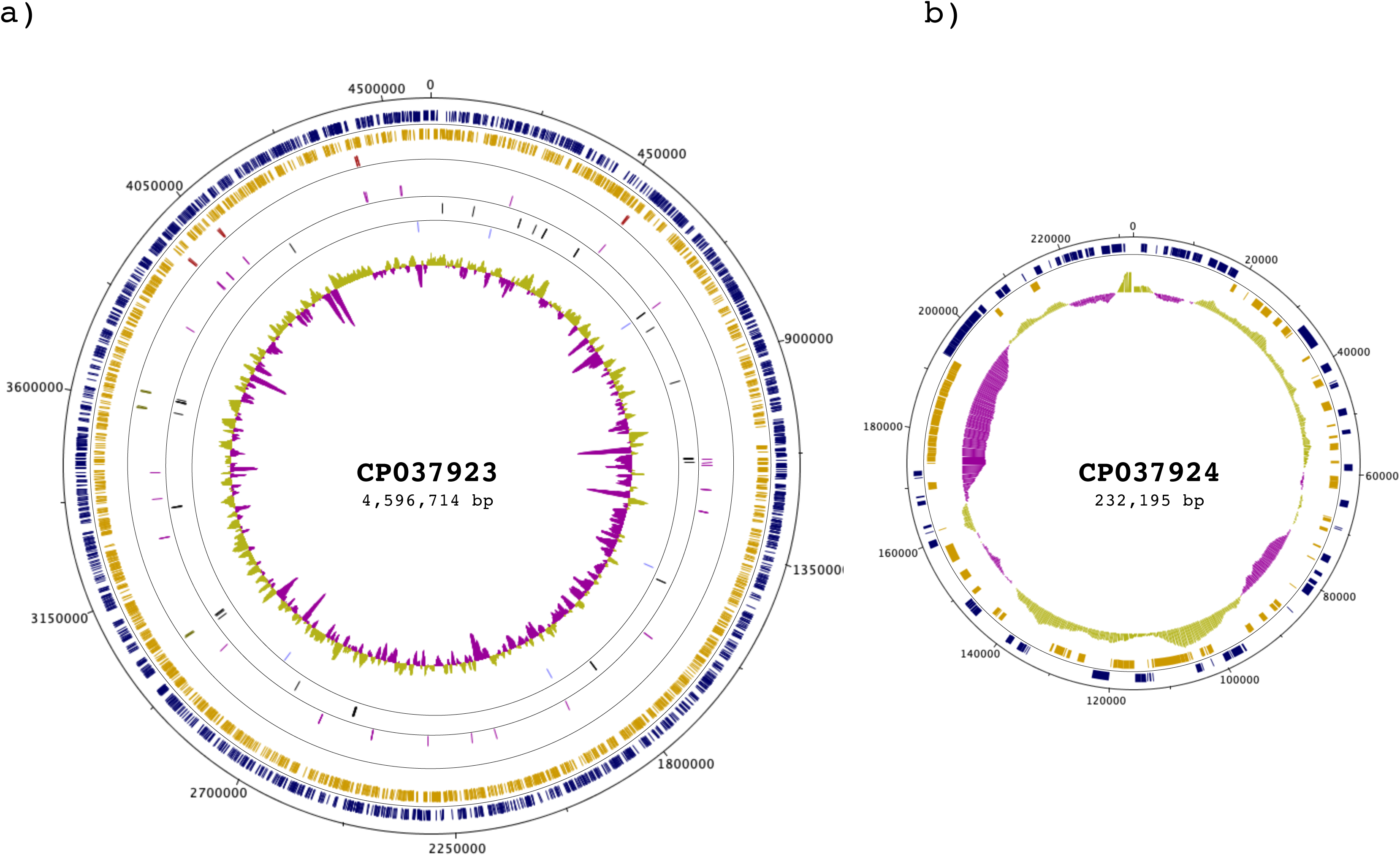
Circular map of the *S. flexneri* 5a M90T genome. The genome is composed of a chromosome (CP037923) and one plasmid (CP037924). The outermost ring represents the nucleotide position (continuous, black). The two following rings within the scale ring depict coding regions (CDSs) in the forward (blue) and reverse (yellow) strand. Moving towards the center, the next rings depict the rRNA in the forward (red) and reverse (green) strand, followed by rings showing the tRNAs in the forward (purple) and reverse (black) strands. The next ring depicts ncRNAs in both strands (light blue). The following ring shows regulatory elements on both strands (light green). The innermost ring shows the GC content. The figure was generated by Circular-Plot[63].

### Polishing of genome assembly using RNA-seq reads

We employed reads from RNA-seq experiments performed on an Illumina HiSeq2000 system to polish the assembled genome. For the first round of polishing, we used the BWA software[26, 27] to align with the assembled genome the reads generated from a library in which the rRNA was depleted with RiboZero (RNAseq-RZ). This step allowed us to polish all the transcribed regions, independently of post-transcriptional processing, as with this method of rRNA depletion all other classes of RNAs are retained. The resulting alignment was used to feed Pilon/1.22[28] for a second round of iterative genome assembly polishing. This second round of polishing was performed with the data set generated with RNA from which the rRNA was depleted with 5’-phosphate-dependent Exonuclease (RNAseq-TEX). The polishing process was stopped when no further changes were observed in the Pilon output. The final polished genome assembly yielded the chromosome, 4,596,714 bp in length (Fig. 1a), and the virulence plasmid pWR100, 232,195 bp in length (Fig. 1b). Both replicons were gap-free and circular molecules.

### Gene prediction and functional annotation

Gene prediction and annotation was carried out with three different pipelines; a) RAST, b) Prokka and c) PGAP/NCBI. The number of predicted genes, coding regions (CDSs) and non-coding RNAs (ncRNAs) was different for the three analyses (Table S1). This can, at least in part, be explained by the fact that all the pipelines used for gene prediction and annotation employ different databases for homology searches. The RAST pipeline used the Taxonomy ID: 1086030 from NCBI, which corresponds to *S. flexneri* serotype 5a strain M90T. RAST predicted and annotated 5,299 genes, but didn’t predict any ncRNA or pseudogene. The pipeline PGAP/NCBI predicted and annotated a lower number of genes, 4,996, of which, 769 are pseudogenes (frameshifted=406, incomplete=305, internal stop=166 and multiple problems=103). From the 769 pseudogenes, 640 were predicted on the chromosome and 129 on the virulence plasmid. The Prokka pipeline predicted and annotated a higher number of genes, 5,021 along with 220 ncRNAs. The most striking feature of Prokka which is distinct from other pipelines is the use of multiple databases to find sequence homologies. For subsequent analysis, we selected the PGAP/NCBI annotation. However, gene annotations with RAST and Prokka are available in the supplementary information in the GenBank format (File S1, S1.1, S2 and S2.1).

Our data showed that *S. flexneri* 5a M90T has a high number of pseudogenes (Table 1) and insertion sequences (ISs) (Table 2). In the genome of *S. flexneri* 5a M90T that we are reporting there are 13 different families of ISs on the chromosome and 15 families on the virulence plasmid (Table 3). Pseudogenes are defined as fragments of once-functional genes that have been silenced by one or more nonsense, frameshift or missense mutation. Pseudogenes can be the result of errors during the replication process or the effect of ISs that shift the open reading frame and modify the DNA sequence. The silencing of the genes can be at two different level, a) Transcriptional or b) Translational. We verified the expression of the identified pseudogenes, both in the chromosome and in the plasmid, using our RNA-seq data (described later). Our results show that 99% of all identified pseudogenes are transcribed, many highly, indicating that their inactivation did not occur at transcriptional level at least (Fig. 2). The *S. flexneri* 5a M90T annotated genome sequence is integrated into RegulonDB[29](http://regulondb.ccg.unam.mx) and RSAT[30] (http://embnet.ccg.unam.mx/rsat/) for use its analysis tools in*S. flexneri* 5a M90T genome.

**Table 1.** General features of the *S. flexneri* 5a M90T genome compared with the sequence and annotation of the previous versions. a) Features in the virulence plasmid, b) features in the chromosome.

|  | Prokka |  | RAST |  | PGAP/NCBI |  |
| --- | --- | --- | --- | --- | --- | --- |
|  | Chromosome | pWR100 | Chromosome | pWR100 | Chromosome | pWR100 |
| **Total length (bp)** | 4596714 | 232195 | 4596714 | 232195 | 4596714 | 232195 |
| **No. total CDSs** | 4713 | 310 | 4943 | 364 | 4629 | 320 |
| **No. Of rRNAs** | 22 | 0 | 22 | 0 | 22 | 0 |
| **No. of tRNAs** | 103 | 0 | 102 | 0 | 102 | 0 |
| **No. of pseudogenes** | 0 | 0 | 0 | 0 | 640 | 129 |

**Table 2.** Insertion sequences (ISs) identified in *S. flexneri* 5a M90T. A) ISs identified on the chromosome, b) ISs identified on the virulence plasmid (pWR100).

| Accession number<br>GenBank | Length (bp) | Genes | CDSs | ISs | Pseudogenes | Reference |
| --- | --- | --- | --- | --- | --- | --- |
| NC_024996 | 213494 | 104 | 104 | 22 | 5 | Buchrieser, C. *et al.*, 2000. |
| AF348706 | 221851 | 294 | 293 | 153 | 0 | Venkatesan. M. M., *et al.*, 2001. |
| CP037924 | 232195 | 307 | 320 | 106 | 129 | This work |

b) Features of the *S. flexneri* 5a M90T chromosome.
| Accession number<br>GenBank | Length (bp) | Genes | CDSs | rRNA | tRNA | ISs | Pseudogenes | Reference |
| --- | --- | --- | --- | --- | --- | --- | --- | --- |
| CM001474 | 4580866 | 4605 | 4013 | 22 | 99 | 385 | 197 | Onodera, N. T. *et al*, 2011. |
| CP037923 | 4596714 | 4049 | 4629 | 22 | 102 | 296 | 640 | This work |

a) Insertion sequences in the *S. flexneri* 5a M90T chromosome.
| Insertion sequence type | No. of IS |
| --- | --- |
| IS1 | 109 |
| IS110 | 6 |
| IS200/IS605 | 3 |
| IS3 | 73 |
| IS3-like | 44 |
| IS4 | 21 |
| IS4-like | 3 |
| IS481 | 1 |
| IS66 | 20 |
| IS66-like | 6 |
| IS91 | 7 |
| ISC | 1 |
| ISNCY | 2 |
| Total | 296 |

b) Insertion sequences in *S. flexneri* 5a M90T pWR100.
| Insertion sequence type | Number of IS |
| --- | --- |
| IS1 | 11 |
| IS110 | 2 |
| IS110-like | 2 |
| IS21 | 2 |
| IS256 | 2 |
| IS3 | 33 |
| IS3-like | 4 |
| IS4 | 5 |
| IS4-like | 1 |
| IS5 | 4 |
| IS630 | 4 |
| IS66 | 21 |
| IS66-like | 3 |
| IS91 | 9 |
| ISL3 | 3 |
| Total | 106 |

**Figure 2.**
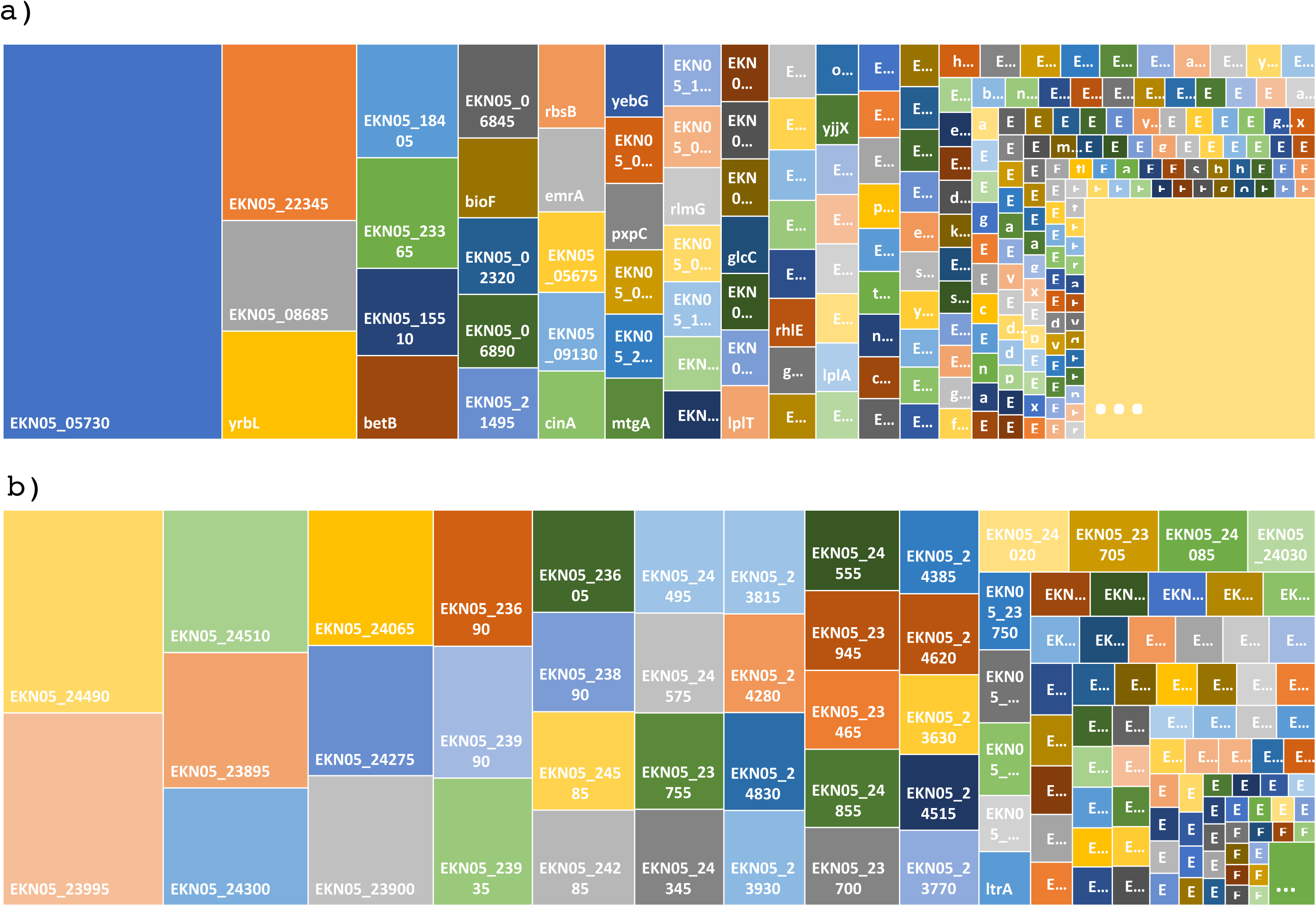
Treemap plot of pseudogenes transcription abundance level in *S. flexneri* 5a M90T. The size of every box is proportional to transcript abundance. RNA-seq total number of reads per gene counted with htseq/0.9.1, a) Pseudogenes transcribed in the chromosome, b) Pseudogenes transcribed in the virulence plasmid pWRl00.

### Whole-genome transcriptional start site determination

To obtain differential RNA-seq (dRNA-seq) data, RNA samples were prepared from triplicate *S. flexneri* 5a M90T cultures grown in TSB at 37 °C and 150 RPM until OD_600_ =0.3. This resulted in a dataset of ∼ 120 million reads mapped to the previously completed reference genome of *S. flexneri* 5a M90T. A total of 14,051 TSSs (Fig. 3) were automatically annotated with ReadXplorer[31] based on the dRNA-seq data and evenly distributed on the forward and the reverse strands. Then, these were categorized according to their position in relation to the annotated genes. TSSs located ≤300 nt upstream of the start codon and on the sense strand of an annotated gene were designated as primary transcriptional start site (pTSS). TSSs within an annotated gene were designated as secondary TSS (sTSS). On the virulence plasmid we annotated 835 TSSs, of which 443 were categorized as primary and 392 as secondary TSSs (Fig. 3). For the chromosome we annotated 13,216 TSSs, of which 6,280 were designated as primary TSSs and 6,936 as secondary TSSs (Fig. 3, Table S2 and S3). In total we have annotated 6,723 putative pTSSs and 7,328 putative sTSSs. This number corresponds to roughly 2.7 TSSs per CDS. The global TSS map of *S. flexneri* 5a M90T and the genome sequence has been integrated into RegulonDB(http://regulondb.ccg.unam.mx/)[29] for easy accessibility and visual display.

**Figure 3.**
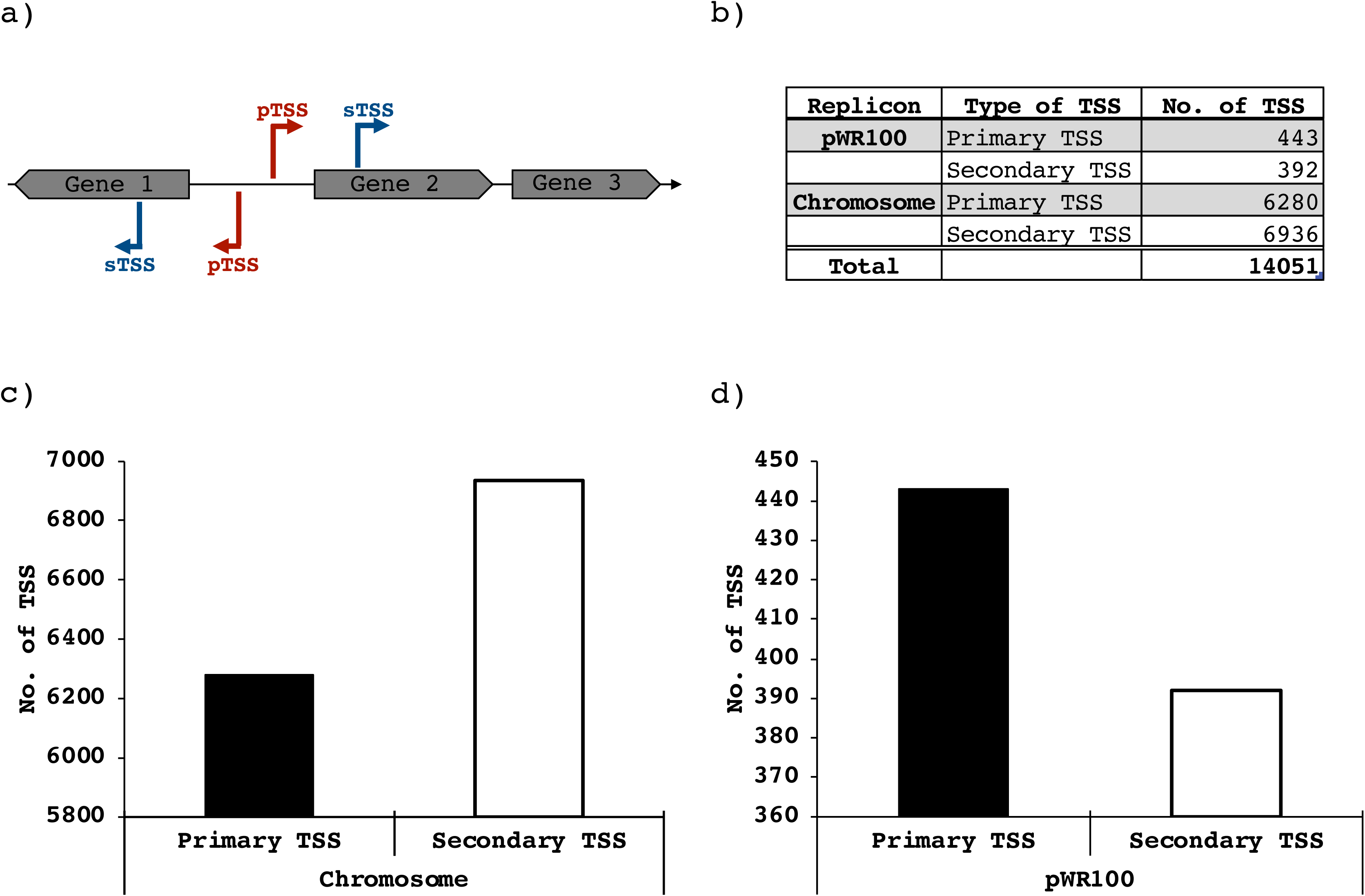
Number of identified transcriptional Start Sites (TSSs) in *S. flexneri* 5a M90T grown in TSB at OD _600_ =0.3. a) Schematic representation of primary TSSs (pTSSs) and secondary (sTSSs), b) Table of TSS by replicon, c) Number of TSS identified on the chromosome, d) Number of TSS identified on the virulence plasmid pWR100.

### Analysis of the length of 5’-UTRs and leaderless transcripts

The TSS analysis shows that the longest 5’UTR in *S. flexneri* 5a M90T is 190 bp on the chromosome and 128 bp on the virulence plasmid (Fig. 4), while the shortest leader in both replicons is only 1 nt long. The average length of leaders on the virulence plasmid is 18 nt and 20 nt on the chromosome. Most primary and secondary TSSs have a 5’-UTR of variable length, but we have found 172 TSSs without leader region on the chromosome and 6 on the virulence plasmid (Table S2 and S3). The graphical visualization of 5’-UTRs is available at RegulonDB (http://regulondb.ccg.unam.mx/).

**Figure 4.**
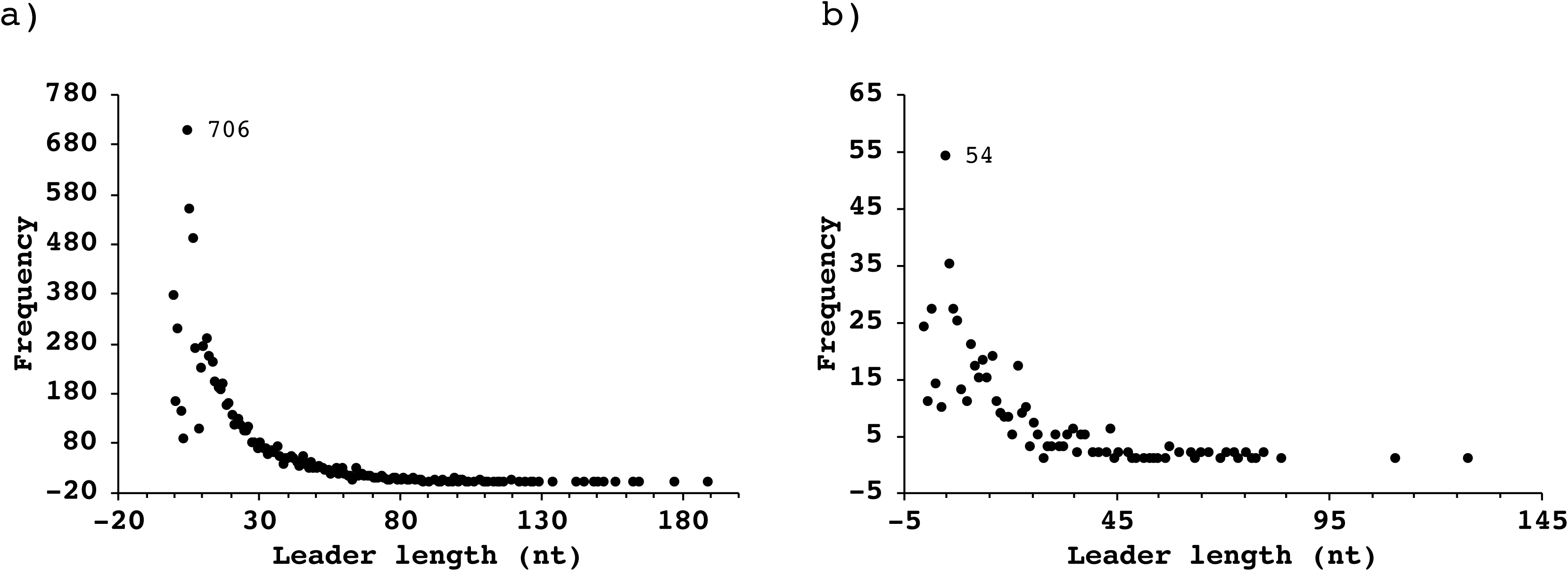
Histogram of 5’-UTR lengths in *S. flexneri* 5a M90T. The distribution of 5’-UTR lengths ranges from O (leaderless) to maximum 190 nt. Transcripts with a 5’-UTR of 5 nt is the most abundant. a) 5’-UTR lengths in the chromosome, b) 5’-UTR lengths in the virulence plasmid.

### Data accessions

The fully sequenced and annotated *S. flexneri* 5a M90T genome is available in GenBank under the accession numbers CP037923 (chromosome) and CP037924 (virulence plasmid). The raw data from PacBio and Illumina sequencing are available in the SRA database under the accession SRR8921221(RNAseq-RiboZero), SRR8921222(dRNA-Seq_TEX_Positive), SRR8921223 (dRNA-Seq_TEX_Negative), SRR8921224(PabBio raw data) and SRR8921225 (RNAseq-TEX). The expression data set is available in GEO[32] with the accession number XXXXX.

As this is the only full genome of *S. flexneri* 5a M90T, it has been recognized as the reference genome and included in the RefSeq database with the accession numbers NZ CP037923 (chromosome) and NZ CP037924 (virulence plasmid). All data that were generated are integrated into RegulonDB for easy accessibility and visualization with JBrowser[33]. The *S. flexneri* 5a M90T genome is integrated in RSAT[30] database to use its analysis tools.

## Discussion

The genome sequence that we report here contains some substantial differences compared to the previously sequenced versions[10, 11, 14]. Minor differences might be due to the fact that the previously published DNA sequences of *S. flexneri* 5a M90T were obtained from a streptomycin-resistant spontaneous mutant *(S. flexneri* 5a M90T Sm), which was derived from the original *S. flexneri* 5a M90T isolate sequenced here by serial culturing on antibiotic-containing plates[14, 34]. Nevertheless, most of the differences can be ascribed to technological developments. On the one hand, the *S. flexneri* 5a M90T chromosome was previously sequenced with a short-read Illumina sequencer[14]. On the other hand, the earlier published virulence plasmid sequences were obtained using short-read ABI377 technology[10, 11]. Repetitive or AT-rich regions make it difficult to prepare a complete genome sequence with short-reads technologies [10, 11]. However, these assembly and annotation problems are circumvented with long-read sequencing such as the PacBio technology employed here. The virulence plasmid described previously[10] is 213,194 bp long (RefSeq accession number NC_024996) [10], 8,357 bp shorter than the sequence published one year later (GenBank accession number AF348706)[11]. The virulence plasmid sequence that we report here is 10,344 bp longer than the one with accession number AF348706[11] (Table 2). It is already known that the genomic structure of the virulence plasmid is a mosaic with many repeated regions and AT-rich tracks[11]. The chromosome that was sequenced previously is 4,589,866 bp in length (GenBank accession number CM001474) including many regions with gaps that are represented in the sequence with “N”. The total number of “N” in the genome scaffold is 11,901 bp[14]. Probably, many of these regions are repeated sequences or AT-rich tracks. The chromosome sequence reported here is gapless, it includes the missed 11,901 bp in the previously reported chromosome sequence[14] plus an extra 15,848 bp that were not present in the previously sequenced scaffold chromosome sequence. All together, these new regions in the genome sequence are summing up to a total 27,749 bp extra on the chromosome.

The number of genes that have been annotated in *S. flexneri* 5a M90T earlier is smaller[14] because of the technical advances in sequencing and computer power for genome analysis and annotation (Table 1). The genome reported here has 4,949 CDSs, of which 4,629 are on the chromosome, in contrast to 4,013 CDSs annotated on the chromosome previously[14] (Table 1), resulting in 616 new CDSs. This discrepancy in the numbers of CDSs could be a consequence of gaps in the previously reported chromosome sequence. In the case of the virulence plasmid, we report 27 new CDSs (Table 1) compared to the last sequence[11].

Of all the annotated genes, 769 are putative pseudogenes (Table 1), which is a high number. When bacteria evolve from free-living to intracellular, the genome is modified. For example, intracellular pathogenic bacteria, such as *Mycobacterium leprae*, may accumulate a large number of pseudogenes [35, 36]. Pseudogenes are particularly prevalent in bacterial species that have recently become associated with or are dependent on eukaryotic hosts, as is the case of *Salmonella*[37]. Pseudogenes are continually created in bacterial genomes from ongoing mutational processes and are subject to degradation and eventual removal by further accumulation of mutations. Besides *Salmonella*, in other members of the *Enterobacteriaceae* such as *E*. *coli* and various *Shigella* species or strains, many CDSs have been annotated as pseudogenes[8, 15, 37–39]. The numbers reported here are in the same range as found in other bacteria such as *Salmonella enterica*[37], *Helicobacter pylori*[16] and *Streptomyces coelicolor*[40]. The process of gene conversion into pseudogenes or gene decay is much faster in *Shigella* than in *E. coli*[41], possibly reflecting a series of key evolutionary events in the conversion of *Shigella* from a commensal to an intracellular pathogen. It has been shown that *Shigella* diverged from its common *E. coli* ancestor in several independent events[42]. Previous studies reported the inactivation of genes that hamper intracellular life in *S. flexneri* 5a M90T by various types of mutations, for example in the *nadA, nadB* locus encoding the capacity to synthetize cadaverine[43, 44] or the *fim* cluster encoding fimbriae[45], pointing towards an adaptative process driving the intracellular style life of *S. flexneri*.

In the present study, we show that the pseudogenes that are present in the genome are transcribed, some of which highly, under laboratory growth conditions (Fig. 2 and Table S4).

Pseudogenes and ISs appear to drive the bacterial genome remodeling. The presence of a large number of pseudogenes in a genome usually correlates with a high number of ISs[46]. The ISs play a very important role, particularly in genome evolution of pathogenic bacteria[47]. ISs transposition can have different outcomes, from simple gene inactivation to constitutive expression or repression of adjacently located genes by delivering IS-specified promoter or terminator sequences, respectively. Multiple copies of ISs elements promote various genomic rearrangements such as inversions, deletions, duplications and fusions between replicons. ISs are determining factors for the efficiency of gene transfer between different bacterial strains or species[47]. Here, we show that the proportion of ISs in the virulence plasmid is much higher than in the chromosome, indicating that the plasmid undergoes more active genome remodeling (Table 2).

Recent evidence indicates that the transcriptional architecture of bacteria is far more complex than originally proposed[48]. The classical model of an operon comprises a group of genes under the control of a regulatory protein, where transcription results in a polycistronic mRNA with a single TSS and a single transcriptional terminator site (TTS). This classical model of operon may be valid for a specific group of genes under specific growth or environmental conditions. However, more recently, many examples have established that an operon may encode alternative transcriptional units which are active under varying environmental conditions[23]. For example, for *E. coli* MG1655, which encodes roughly 4600 genes, >14,000 TSSs were documented[49, 50]. Our results for *S. flexneri* 5a M90T in terms of TSSs number – about 14,051 TSSs for a genome of 4.8 Mb - are close to what was found in *E*. *coli* MG1655.

To achieve a complex landscape of alternative transcriptional units, transcriptional regulation occurs at multiple levels[51]. Different lengths of the 5’-UTR play a very important role in translational regulation[52–54]. Indeed, the length of a 5’-UTR can provide insight into the regulation of gene expression[53]. For example, long 5’UTRs may contain riboswitches or provide binding sites for small regulatory RNAs[55]. Leaderless genes are translated by a different mechanism than genes with a leader sequence, as is the case for *virF* in *Shigella* spp.[56].

## Conclusions

The genome sequence reported here is the first complete, gapless genome sequence for *S. flexneri* 5a M90T. Automatic annotation combined with manual curation allowed us to provide a high quality reference genome that will be extremely useful to several types of studies, for example transcriptomics, differential expression analyses, or genome evolution. Moreover, in molecular pathogenesis projects, our results can be used as a resource to know which genes are transcribed before infection of host cells. The genome sequence together with the analysis of transcriptional start sites is also a valuable tool for precise genetic manipulation of *S. flexneri* 5a M90T.

In the present work we describe a hybrid cutting-edge sequencing approach and produced a high-quality, gapless reference genome. The hybrid pipeline that we report here for genome sequencing with long reads and polishing with RNAseq data proofed as a powerful strategy for genome assembly, polishing and annotation that could be implemented any type of organism.

*S. flexneri* serotype 5a strain M90T is a very important model to study the molecular pathogenesis mechanisms. The availability of a full genome opens the door to discovering new genomic elements and gene regulatory networks that are involved in *Shigella* pathogenicity.

## Material and methods

### Bacterial strain and culture conditions

The *S*. flexneri serotype 5a strain M90T that was used in this study was obtained from Dr. Philippe Sansonetti, Institut Pasteur, Paris, France. This strain was collected by Dr. Samuel Formal for the Walter Reed Army Institute of Research collection and first described in 1982[9]. This strain is not streptomycin resistant, unlike a derivative of M90T that was later obtained by serial passaging on antibiotic containing plates[34] and sequenced in 2012[14]. *S. flexneri* 5a M90T was cultured on tryptic soy broth agar plates with 0.01% (w/v) Congo red (TSBA-CR). Red colonies were selected to ensure the presence of the virulence plasmid (pWR100). Overnight bacterial cultures were grown at 37°C in tryptic soy broth medium, sub-cultured 1:100, and grown at 37°C in a shaking incubator at 150 RPM, until OD_6oo_=0.3.

### DNA purification and genome sequencing

Genomic DNA was isolated using the WizardR Genomic DNA purification kit (Promega, Inc.) from overnight cultures of *S. flexneri* 5a M90T according to the manufacturer’s instructions. Isolated DNA was cleaned as many times as necessary with phenol-chloroform (until no white interphase between the watery and organic phase was forming)[57] to obtain a high quality-quantity of genomic DNA (20 µg) for PacBio library preparation[12]. Library preparation was carried out by Novogene Inc. Sequencing was performed using a PacBio RSII sequencer at Novogene Inc, Hong Kong, China.

### Total RNA purification and sequencing

*S. flexneri* 5a M90T was sub-cultured until OD_600_ =0.3 and the culture was mixed with 0.2 volumes of stop solution (95% EtOH and 5% phenol pH 4, v/v)[57]. Samples were allowed to incubate on ice for at least 30 min, but not longer than 2 h, to stabilize the RNA and prevent degradation. After the incubation on ice, the cells were harvested by centrifugation for 5 min at 13,000 RPM at 4°C. Cell pellets were frozen with liquid nitrogen and stored at −80°c until RNA extraction.

Frozen cell pellets were thawed on ice and resuspended in lysis solution (0.5% SDS, 20 mM sodium acetate pH 4.8, 10 mM EDTA pH 8). Bacterial cells were lysed by incubating the samples for 5 min at 65°C. Afterwards, total RNA was extracted using the hot-phenol method[16]. Contaminating DNA was digested by DNase I (Roche; 1 U/µg RNA, 60 min, 37°C) in the presence of RNase inhibitor (RNaseOUT, ThermoFisher Scientific; 0.1 U/µl) followed by clean-up of RNA by phenol/chloroform/isoamyl alcohol and precipitation of RNA with 2.5 volumes of ethanol containing 0.1 M sodium acetate pH 5.5 and 20 µg of glycogen (Roche)[57]. Removal of residual DNA was subsequently verified by control PCR using the oligos SF-Hfq-F 5’-ACGATGAAATGGTTTATCGAG-3’ and SF-Hfq-R 5’-ACTGCTTTACCTTCACCTACA-3’, which amplify a 909 bp long product of the *hfq* gene from *S. flexneri* 5a M90T including 300 pb upstream and 300 bp downstream of the *hfq* gene.

The RNA concentration was measured using a NanoDrop ND-1000 spectrophotometer (Saveen & Werner AB, Limhamn, Sweden). Thereafter, the integrity of the 16S and 23S rRNA was checked by agarose gel electrophoresis, using 1% agarose in 1X TAE buffer (40 mM Tris acetate, 1 mM EDTA at pH 8.3±0.1).

The rRNA was depleted from three biological replicates of total RNA with RiboZero according to the manufacturer’s instructions (Illumina, Inc.). Library preparation and sequencing were performed at the EMBL Genomics Core Facility (Heidelberg, Germany).

The rRNA from another set of three biological replicates was depleted with Terminator-5’-Phosphate-Dependent Exonuclease (TEX) (Lucigene, Inc.). Library preparation and sequencing was performed at Novogene, Inc. The libraries were constructed using the TruSeq Stranded kit and were sequenced on an Illumina HiSeq2000 platform (Illumina, Inc.) with a paired-end protocol and read length of 150 nt (PE150), resulting in a total output of roughly 20 million (M) per sample. All reads outputs were checked for passage of Illumina quality standards[58, 59]. These RNA-seq results obtained from EMBL and Novogene were used to polish the genome assembly.

### Genome assembly and annotation

*De nova* genome assembly was performed with the script Canu/1.7[25] implementing the pacbio-raw option using all its default parameters. Output files from Canu assembly were used as input to polish the genome assembled with Pilon/1.22[28]. Polishing of the genome assembly was done in two rounds: first one was carried out using the RNA-seq output files from the samples in which the rRNA was depleted with RiboZero (RNAseq RZ) and the second one was carried out with the RNA-seq results from the samples in which the rRNA was depleted with TEX (RNAseq-TEX). Genome annotation and polishing was performed at Uppsala Multidisciplinary Center for Advanced Computational Science (UPPMAX) of SciLifeLab at Uppsala University, Sweden.

Annotation of assembled contigs was done using three different pipelines: 1) NCBI Prokaryotic Genome Annotation Pipeline (PGAP)[60], 2) Prokka/1.12-12547ca[61] and 3) Rapid Annotation using System Technology version 2.0 (RAST)[62]. Genome annotation with Prokka was performed on the UPPMAX server facilities at Uppsala University, Sweden.

The assembled and annotated genome was manually curated using Artemis[63] for visualizing and editing the genome files. The genome was deposited in GenBank with accession numbers CP037923 (chromosome) and CP037924 (virulence plasmid).

### RNA treatment for transcriptional start site (TSS) determination and sequencing

To determine transcriptional start sites, the RNA of three biological replicates in which the rRNA had been depleted with RiboZero (Illumina, Inc.) was used. To enrich for primary transcripts, we exploited the property that primary bacterial transcripts are protected from exonucleolytic degradation by their triphosphate (5’PPP) RNA ends[24], while RNAs containing a 5’ monophosphate (5’P) are selectively degraded[16, 24]. The rRNA-depleted RNA was split into two aliquots. One aliquot was treated with Terminator 5’-Phosphate-Dependent Exonuclease (TEX+), the other aliquot was incubated only with TEX buffer (TEX-) as a control. TEX treatment was carried out for 60 min at 30°c. One unit of TEX was used per 1 µg of rRNA-depleted RNA. Following organic extraction (25:24:1 v/v phenol/chloroform/isoamyl alcohol), RNA was precipitated overnight with 2.5 volumes of ethanol/0.1M sodium acetate (pH 5.5) and 20 µg of glycogen (Roche) mixture. After TEX treatment, both samples (TEX+ and TEX-) were treated with 5’, Pyrophosphohydrolase (RppH) (NewEngland BioLabs, Inc.) to generate 5’-mono-phosphates for linker ligation[64], and again purified by organic extraction and ethanol precipitation. RppH treatment was carried out for 60 min at 37°c. An RNA adaptor (5’ GACCUUGGCUGUCACUCA-3’) was ligated to the 5’-monophosphate of the RNA end by incubation with T4 RNA ligase (NewEngland BioLabs, Inc.), at 25°c for 16 h. As last step, the RNA adaptor that had been ligated to the RNA was phosphorylated with T4 PNK (NewEngland BioLabs, Inc.) at 37°C for 60 min.

Separate libraries were constructed for TEX- and TEX+ samples. The libraries were constructed using TruSeq Stranded Kit (Illumina, Inc.) and sequenced on an Illumina HiSeq2000 platform (Novogene, Inc.) with a paired-end protocol and read length of 150 nt (PE150), resulting in a total output of roughly 20 million (M) per sample/library sequenced. All reads were checked for passage of Illumina quality standards[58, 59].

### Reads mapping of TSS library

Reads in the FASTQ format were cleaned up with trimmomatic/0.36[65] to remove sequences originating from Illumina adaptors and low quality reads. The files were aligned with the previously assembled reference genome of *S. flexneri* 5a M90T (GenBank accession numbers CP037923 and CP037924) with bowtie2/2.3.4.3[66] using −X 1000 such that only mate pairs were reported if separated by less than 1000 bp. All the other settings were implemented with the default option. After the alignment was completed, samtools /1.9[26] was used to remove duplicates and select for reads that were aligned in proper pairs. Reads aligned to the reference genome was converted to BAM format with samtools/1.9. The final analysis for identification and annotation of TSSs into pTSS and sTSS was done with ReadXplorer[31, 67].

### Reads mapping of transcribed pseudogenes

To count the number of reads aligned per pseudogene we used the alignment results for TSSs mapping. After sorting the alignment with samtools/1.9[26], the reads counting per pseudogene was performed with htseq/0.9.1[68] using stranded mode and pseudogene as a feature type. The total expression of pseudogenes is the average of the six libraries used for TSSs determination.

### Transcriptional start sites annotation and classification

To map dRNA-seq outputs, reads were split by replicon, converted to BAM format and sorted by position with samtools/1.9[28]. These BAM files were used as input for ReadXplorer[31, 67] for automatic *de novo* TSSs annotation. For the analysis, results from three biological replicates were pooled and TSS within 10 nt of each other were clustered into one. Such regions were then manually annotated by scanning the respective wiggle files for nucleotides with an abrupt increase in coverage. Transcriptional start sites were classified according to their genomic context. Peaks in an intergenic region and on the same strand as the closest downstream gene were classified as primary. Peaks within gene boundaries and on the same strand as the gene were qualified as secondary. All TSS positions were assigned relative to the start of the associated annotated gene. With the first base of the gene being positive +1, all upstream position start with - 1.

## List of abbreviations

IS: Insertion sequence
SMRT: Single molecule real-time
TSS: Transcriptional start site
TEX: 5’-monophosphate-dependent exonuclease
TSB: tryptic soy broth
CDS: Coding sequence
ncRNA: Non-coding RNAs
dRNA-seq: Diffential RNA sequencing
pTSS: Primary transcriptional start site
sTSS: Secondary transcriptional star site
TTS: Transcriptional terminator site
TSBA-CR: tryptic soy broth with 0.01% (w/v) Congo red.

## Declarations

### Ethics approval and consent to participate

Not applicable.

### Consent for publication

Not applicable.

### Availability of data and material

Data are deposited at GenBank database under the accession number: CP037923 and CP037924.

The raw data are deposited in SRA under the accession number SRR8921221(RNAseq-RiboZero), SRR8921222(dRNA-Seq_TEX_Positive), SRR8921223 (dRNA-Seq_TEX_Negative), SRR8921224(PabBio raw data) and SRR8921225 (RNAseq-TEX). https://dataview.ncbi.nlm.nih.gov/

Request for material should be directed to and will be fulfilled by Andrea Puhar.

### Author’s contribution

RCR and ST performed experiments. RCR analyzed all data. RCR and AP designed research. RCR and AP wrote the manuscript. All authors corrected and approved the manuscript.

## Acknowledgements

We are grateful for travel support and courses organized by The National Doctoral Program in Infections and Antibiotics (NDPIA). The computations were performed on resources provided by SNIC through Uppsala Multidisciplinary Center for Advanced Computational Science (UPPMAX) under Project SNIC 2017-7-258.

We are grateful to EMBL Genomics Core Facility (Heidelberg, Germany) for RiboZero library preparation and sequencing.

The authors thank RegulonDB(CCG-UNAM) staff specially to Jaír s. García-Sotelo (LIIGH-UNAM) for their support in data managing and display in RegulonDB, and also RSAT staff to integrate *S. flexneri* 5a M90T genome (CP037923, CP037924) in its analysis tools. We also thank to Dorothee Langenbach, Atin Sharma, David Cisneros (MIMS, Umeå Unversity, Sweden) and Edgardo Sepúlveda (CICESE, Baja California, Mexico) for critically reading the manuscript.

---

**Table S1**. Comparison of general features of the *S. flexneri* 5a M90T predicted with three different pipelines: Prokka[61], RAST[62] and PGAP/NCBI[60].

**Table S2**. Transcriptional start sites determined in *S. flexneri* 5a M90T chromosome with ReadExplorer[31, 67].

**Table S3**. Transcriptional start sites determined in *S. flexneri* 5a M90T virulence plasmid (pWR100) with ReadExplorer[31, 67].

**Table S4**. Pseudogenes transcription abundance level. Reads counting per pseudogene was performed with htseq/0.9.1[68] using stranded mode and pseudogene as a feature type. The total expression of pseudogenes is the average of the six libraries used for TSSs determination.

**File S1**. General features of the *S. flexneri* 5a M90T chromosome annotated with Prokka[61].

**File S1**.**1**. General features of the *S. flexneri* 5a M90T virulence plasmid (pWR100) annotated with Prokka[61].

**File S2**. General features of the *S. flexneri* 5a M90T chromosome annotated with RAST[62].

**File S2**.**1**. General features of the *S. flexneri* 5a M90T virulence plasmid (pWR100) annotated with RAST[62].

